# Detection and discrimination of single nucleotide polymorphisms by quantification of CRISPR-Cas catalytic efficiency

**DOI:** 10.1101/2022.04.22.489229

**Authors:** Charles Blanluet, Diego A. Huyke, Ashwin Ramachandran, Alexandre S. Avaro, Juan G. Santiago

## Abstract

The specificity of CRISPR-Cas12 assays is attractive for the detection of single nucleotide polymorphisms (SNPs) implicated in, *e.g*., SARS-CoV-2 variants. Such assays often employ endpoint measurements of SNP or wild type (WT) activated Cas12 *trans*-cleavage activity; however, the fundamental kinetic effects of SNP versus WT activation remain unknown. We here show that endpoint-based assays are limited by arbitrary experimental choices (like used reporter concentration and assay duration) and work best for known target concentrations. More importantly, we show that SNP (versus WT) activation results in measurable shifts in the Cas12 *trans*-cleavage substrate affinity (*K_M_*) and apparent catalytic efficiency 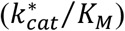. To address endpoint-based assay limitations, we then develop an assay based on the quantification of Michalis-Menten parameters and apply this assay to a 20-base pair WT target of the SARS-CoV-2 E gene. We find that the 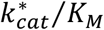 measured for WT is 130-fold greater than the lowest 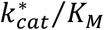 among all 60 measured SNPs (compared to a 4.8-fold for endpoint fluorescence of the same SNP). *K_M_* also offers strong ability to distinguish SNPs, varies 27-fold over all the cases, and is insensitive to target concentration. Lastly, we point out trends among kinetic rates and SNP base and location within the CRISPR-Cas12 targeted region.

## 1 INTRODUCTION

SNP identification has gained critical importance in a range of applications which include genotyping, cancer screening, and COVID-19 diagnostics.^1,2^ The latter specifically involves differentiation of SARS-CoV-2 variants that most often differ from the original sequence by just a few nucleotides.^3^ As a potential solution, CRISPR-based assays with high specificity and point-of-care compatibility have recently emerged.^4^ These assays leverage programmable subtypes of CRISPR associated (Cas) enzymes to identify viral RNA in samples.^5–8^ The capability of Cas enzymes to discriminate among small mutations^7,9–12^ may also be critical to the identification of new SARS-CoV-2 variants.

There have been several reports of CRISPR detection of SNPs in the literature. In particular, Cas12a,^10,13^ Cas13a,^7,9,11,12,14^ and Cas14a^15^ have been used to confirm that CRISPR-based assays are sufficiently specific for SNP detection. For example, Shan *et al*.^14^ showed differences in initial reaction velocities for Cas13 when activated by WT versus SNPs within 6 bases of the target 5’ end but did not consider effects of SNPs on Michaelis-Menten kinetics.

Differentiation among mutants, however, has most often been limited to the use of endpoint fluorescent signal assays which quantify the integral sum of enzyme activity over some specified time. The detailed effects of target mutations on the details of Cas enzyme *trans-cleavage* kinetic rates have not been elucidated. To our knowledge, there has been no systematic study of single base-pair mutations on the Michaelis-Menten kinetics of CRISPR enzymes.

The current work systematically explores the effects of SNPs on LbCas12a enzyme *trans*-cleavage activity. To this end, we use a guide RNA (gRNA) to target a WT, 20-base pair DNA sequence as well as all 60 related SNPs for that sequence. The WT target chosen here corresponds to the E gene of SARS-CoV-2, which is one of the most targeted genetic markers for COVID-19.^16^ In this work, we first demonstrate CRISPR assay conditions wherein endpoint detection of WT and SNP targets appears indistinguishable. We then use a Michaelis-Menten model to measure the Michaelis constant *K_M_* and apparent catalytic efficiency 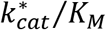 of WT- and SNP-activated Cas12a enzymes. Importantly, we find that *K_M_* and 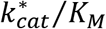 offer effective ability to discriminate WT from mutants in a manner superior to discrimination based on endpoint detection. 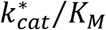 offers the clearest discrimination between the WT and mutants among the methods we measured. However, we hypothesize that discrimination based on *K_M_* alone is itself a powerful assay approach since its quantification does not require knowledge of target concentrations. The current extensive kinetic rate data also exhibits interesting trends among kinetic rates and SNP base type and location. Overall, this study demonstrates SNP effects on the affinity and catalytic efficiencies of *trans*-cleavage and offer improved specificity applicable to CRISPR diagnostics.

## 2 RESULTS

### 2.1 Fluorescence-based detection of SNPs

To assess the ability of endpoint fluorescence assays to discriminate SNPs, we first measured *trans*-cleavage progress curves for the WT and each of the 60 target SNP species (see **Methods** for complete experimental protocol). We therefore complexed LbCas12a with a gRNA and then introduced a complementary WT (or almost complementary SNP ssDNA) target to activate the complexes (**Fig. 1a**). As a control, we used the same concentration of WT and mutant target across all experiments. The incubation of the Cas-gRNA complex with WT and SNP targets initiated *trans*-cleavage of fluorescently labeled ssDNA reporters (**Fig. 1b**). We denote the SNPs using a three-parameter code (*e.g*., T2G indicates that the nucleotide position #2 contains the mutation wherein thymine (T) is replaced by guanine (G)). Heat maps of endpoint fluorescence at, for example, 10 and 45 min (**Fig. 1c**) revealed strong differences in the *trans*-cleavage activity among WT and SNPs. In all heat maps shown in this work, the WT is indicated with an asterisk in the top right corner of the corresponding cell.

**Fig. 1.**
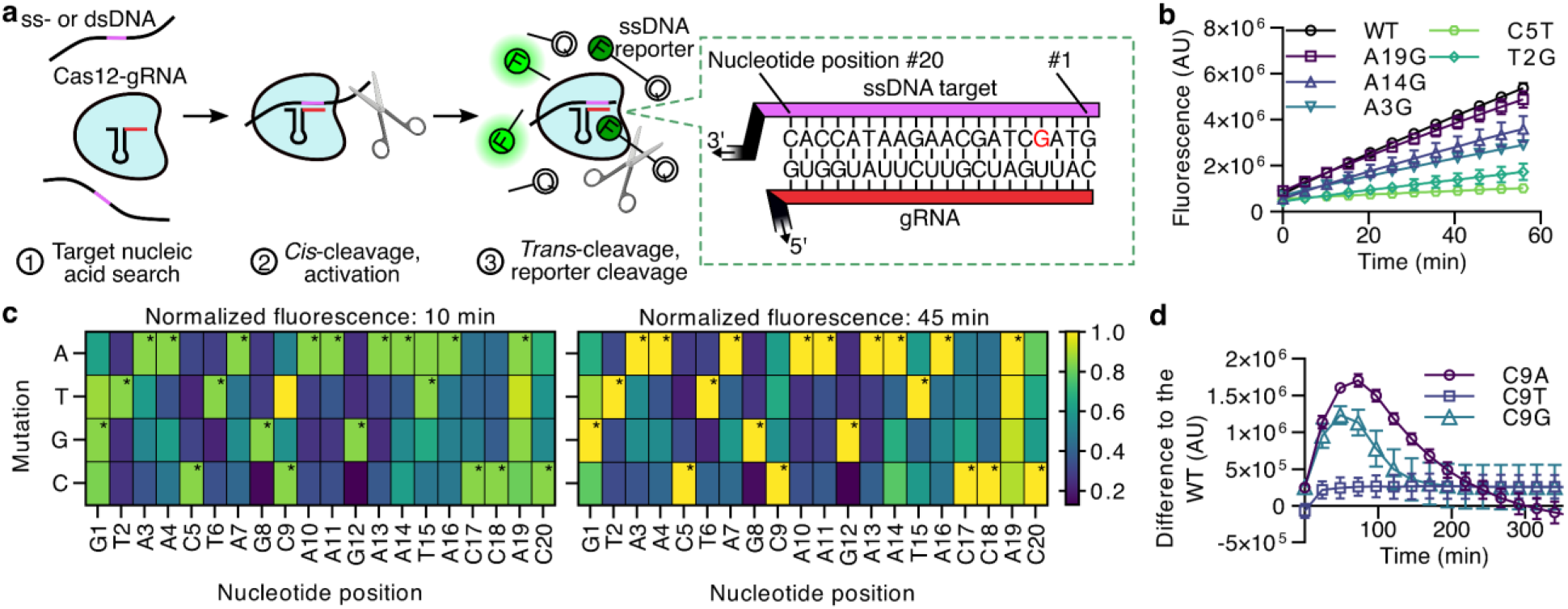
Schematic of CRISP-Cas12a detection and experimental study of endpoint fluorescence detection. **a** Schematic representation of a Cas12-gRNA complex which activates upon recognition of a ssDNA target and initiates *trans*- (or collateral) cleavage of fluorescently labeled ssDNA reporter molecules. The schematic within the green dashed box depicts the interaction between the gRNA and a SNP at position #4 (where a G substitutes an A). **b** Example measured fluorescence versus time for WT and several SNP targets. The Cas12-gRNA complex and reporter concentrations were here 2 and 800 nM, respectively. **c** Fluorescence of WT and SNP targets at 10 min (left column) and 45 min (right column). Data has been normalized to the highest value of each instance. Asterisks denote the WT cells. The enzyme and reporter concentrations match those in **b**. Note C9T is brighter than WT at 10 min. **d** WT fluorescence signal minus that of SNP versus time for SNPs at position #9. The Cas12-gRNA complex and reporter concentrations were respectively 10 and 800 nM.

These experiments show that the degree to which endpoint fluorescence measurements enable discrimination of mutants depends strongly on the time at which the fluorescence was measured. For example, mutant C9T, showed greater fluorescence than WT at 10 min (**Fig. 1c**, left heat map), but significantly less fluorescence than WT at 45 min (**Fig. 1c**, right heat map). To better understand this behavior, we measured the difference in fluorescence signal between WT and SNPs in position #9 (C9T, C9A, and C9G) for longer times and at 5-fold higher target concentrations (**Fig. 1d**). We found that, under our experimental conditions, C9T initially cleaves faster than WT (i.e., its value in **Fig. 1d** is initially negative) but the C9T signal was then overtaken by that of the WT after about 5 min. This behavior is consistent with and can be explained by the underlying kinetics. We demonstrate this using our experimentally validated Michaelis-Menten model^17^ informed by values of *K_M_* and 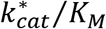 measured here (see **Fig. S1** and discussion). Also, as expected, the WT-to-mutant fluorescence difference for all mutants fluctuated but then plateaued near zero.

These results suggest that endpoint fluorescence is not well suited to differentiate WT from SNPs primarily due to three reasons. First, endpoint fluorescence values depend significantly on the differing amounts of (unmeasured) enzyme activity prior to fluorescence measurements (such as during pipetting or loading). Second, WT-to-SNP endpoint fluorescence signal differences necessarily grow from zero and then asymptote to approximately zero as all reporters are cleaved. Differential *trans*-cleavage rates therefore mean that the choice of measurement time for endpoint detection is critical and the *a priori* choice of reporter concentration strongly influences WT-to-SNP signal differences. Third, the rate of fluorescence increase is largely determined by the (often unknown) concentration of activated enzymes. Hence, a SNP which exhibits a low *trans*-cleavage rate could nevertheless lead to larger fluorescence increase (relative to WT) when the SNP target is in sufficiently higher concentration. As we will discuss, these limitations of endpoint-based detection can be mitigated by characterization of the underlying Michaelis-Menten kinetics. Namely, quantification of *K_M_* and 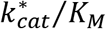 provides a framework to address irregularities during sample preparation, choice of assay time, and unknown target concentrations. Further, quantification of *K_M_* and 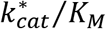 results in significantly greater specificity of detection.

### 2.2 Effect of SNPs on apparent enzyme turnover rate

To characterize the underlying Michaelis-Menten kinetics of SNP-activated *trans*-cleavage activity, we first measured apparent turnover rates 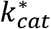. This 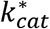 is defined as the ratio of the maximum achievable reaction velocity divided by the target concentration (see **Methods**). That is, 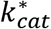 is the value obtained from Michaelis-Menten by assuming that all targets (WT or SNP) result in activated Cas12-gRNA enzyme which are available for *trans*-cleavage kinetics. We report 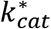 (rather the underlying turnover rate *k_cat_*) since only some (unknown) fraction of these mutants results in activated Cas12-gRNA complexes.^18^ As a systematic comparison, we fixed the Cas12-gRNA complex and target concentrations across experiments to quantify comparable values of 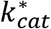. To this end, we first measured *trans*-cleavage progress curves for WT and SNP targets with for reporter concentrations varied between 13 and 1600 nM (example plots for position #4 are shown in **Fig. 2a**). The initial reaction velocities for each reactant system were then fit to a Michaelis-Menten curve (**Figs. 2b** and **S2-S5**). The Michaelis-Menten curve, importantly, yielded estimates of the target-associated 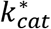 for these and all other SNPs studied here (**Figs. 2c**).

**Fig. 2.**
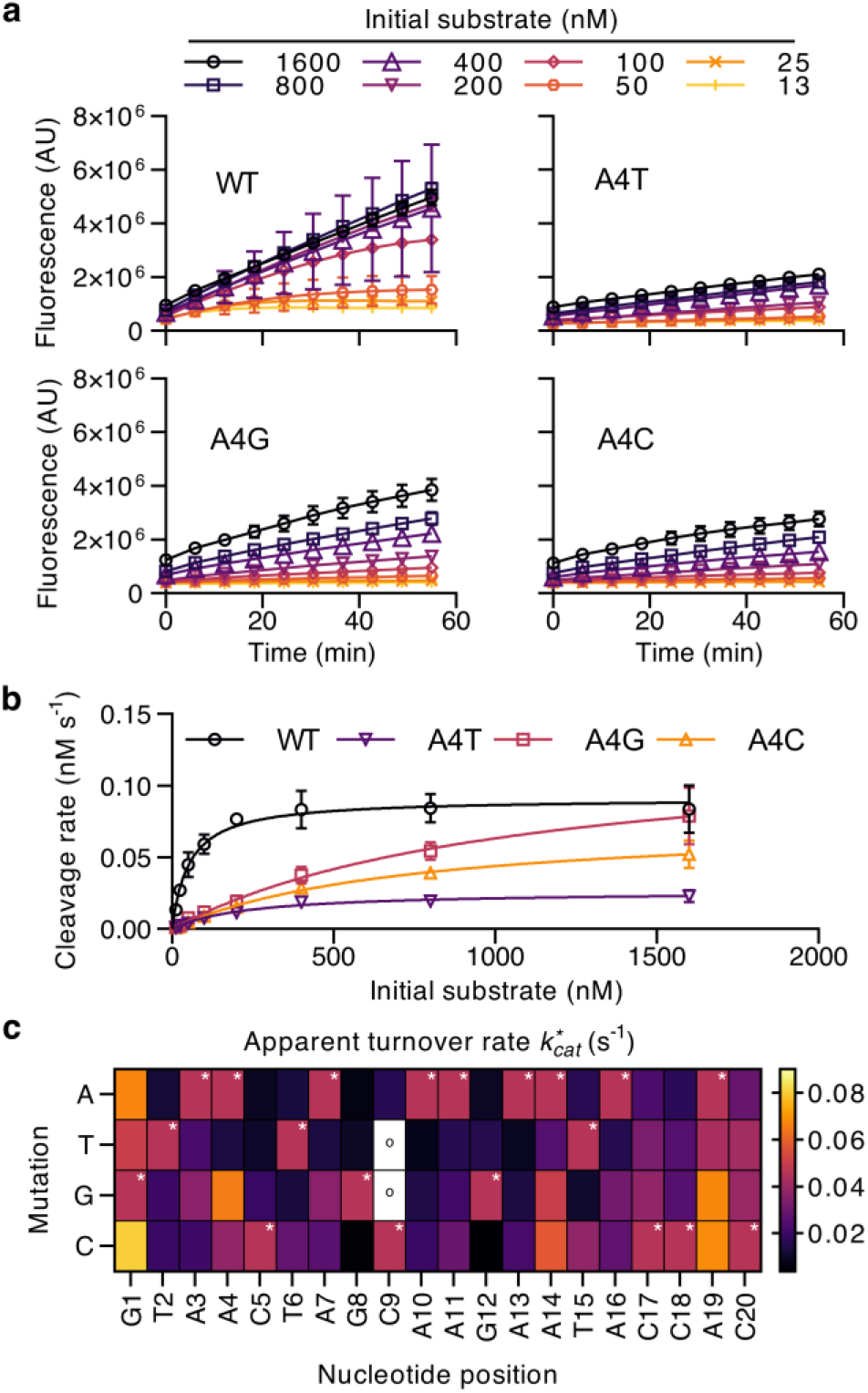
Michaelis-Menten experiments of SNPs and 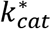 values. **a** Fluorescence versus time curves show *trans*-cleavage of fluorescently labeled ssDNA reporters by a Cas12-gRNA complex activated by WT or SNPs in nucleotide position #4 (all other SNP data is provided in the **Supplementary Information**). **b** Cleavage rate versus initial substrate (uncleaved reporter) concentration for WT and SNPs at position #4. Also shown by the solid lines are fits to the Michaelis-Menten equation. The Cas12-gRNA complex concentration in **a** and **b** was 2 nM. **c** Heatmap of apparent turnover rate 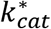 for WT and all associated SNPs, estimated from analysis of the Michaelis-Menten curves. Asterisks denote the WT cells. A small circle denotes cells with 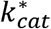 greater than 0.09 s^-1^ (exact values in **Table S1**).

We discovered that 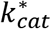 varied widely over the SNPs. For example, the WT 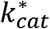 (= 0.046 s^-1^) was surprisingly lower than that of 8 of the 60 SNPs (G1A, G1C, A4G, C9T, C9G, A14T, A19G, and A19C). In fact, the largest 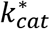 was observed for C9G (4.9-fold greater than the WT 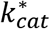). These results suggest that single nucleotide mutations can significantly affect the *trans*-cleavage behavior of Cas12 enzymes in a manner which can, somewhat surprisingly, result in either increases or decreases in enzyme turnover rates. For differentiation among WT and SNP targets, measurements of 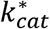 offers a technique that is insensitive to the duration of the experiment (since 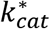 is a fundamental property of the reactant system). Unfortunately, 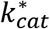 typically requires *a priori* knowledge of the activated enzyme concentration. The latter can be deduced from the Cas12-gRNA complex concentration or the concentration of the target (whichever is lower and therefore limiting). This type of knowledge is, of course, often not realistic in diagnostics of actual biological samples.

### 2.3 Effect of SNPs on Michaelis constant

To detect WT in a manner independent of experiment duration and *a priori* knowledge of target concentration, we turned to the Michaelis constant *K_M_. K_M_* describes the fundamental affinity of the enzyme for substrates is a property unique to a specific combination of Cas12-gRNA, target, buffer chemistry, and temperature. We found that *K_M_* also varied widely across SNPs (**Figs. 3** and **S3-S5**). However, importantly, we discovered that *K_M_* offered much improved discrimination of SNPs versus WT. All but one SNP studied here (C17A, = 43 nM) had higher *K_M_* values (lower affinity) than the WT (= 53 nM). These two values were also well within measurement uncertainty (**Table S1**). Interestingly, when Cas12-gRNA complexes reacted with a mix of WT and SNP targets, almost all complexes were *trans*-activated by WT rather than mutant targets (likely due to the higher affinity of WT targets, **Fig. S6**). We also found that the nucleotide positions nearer to the 3’ end (positions #18 to 20) had *K_M_* values closer to the WT, which suggests a deterministic relation between SNP position and affinity. Interestingly, 6 of the 8 SNPs with a higher 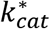 than WT, showed *K_M_* above 500 nM (almost 10-fold higher than WT). Further, the highest *K_M_* was found for SNP C9G (= 1200 nM).

**Fig. 3.**
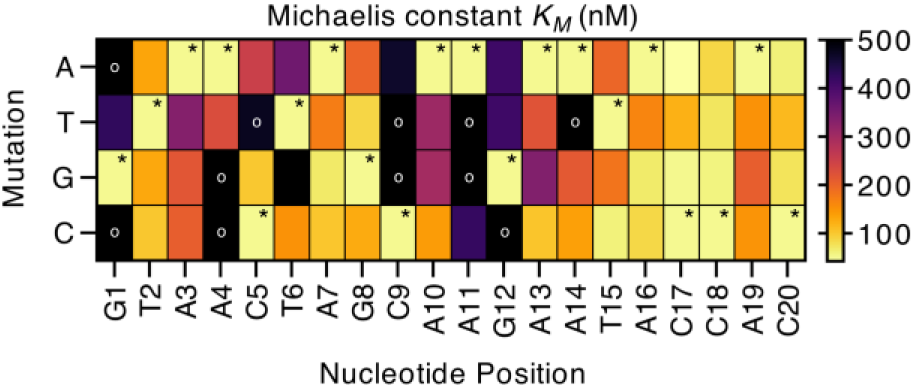
WT differentiation based on Michaelis constant *K_M_*. Heatmap of *K_M_* for WT and all associated SNPs as estimated from analysis of the Michaelis-Menten curve. Asterisks denote the WT cells. A small circle denotes cells with *K_M_* greater than 500 nM (very low affinity, exact values in **Table S1**). We found that all but one of 60 SNPs (C17A) exhibited greater *K_M_* than for WT. This is important as *K_M_* may provide a technique to discriminate WT from SNP that is insensitive to assay time and to target concentration.

The *K_M_* study showed that the WT-activated enzyme complex has a much stronger affinity to the *trans*-cleavage substrates (ssDNA reporters) compared to its SNP-activated counterparts. This strongly suggests that *K_M_* can be a robust method to distinguish WT from SNPs. Most striking is that *K_M_* is a fundamental property of the reactant system which enables WT differentiation without the requirement of a specified assay time or knowledge of target concentration. Instead, the *K_M_* assay directly measures the affinity of differently activated enzymes (WT or mutant) to substrates.

## 3. DISCUSSION

We here discuss the implications of our findings toward the design of highly specific CRISPR-based, *trans*-cleavage assays for SNP detection. Following the determination of 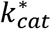 and *K_M_* (both parameters derived from the same experiment), we calculated the apparent catalytic efficiency 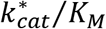. A comparison of all 60 SNPs to the WT shows that 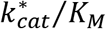 provides highly specific recognition of WT target (**Fig. 4a**). In fact, none of the 60 SNPs yielded a higher 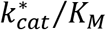 than the WT. The WT 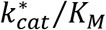 (= 8.6 × 10^5^ M^-1^ s^-1^) was 130-fold higher than the lowest SNP 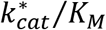. The latter can be compared to the modest 4.8-fold difference observed for endpoint fluorescence of the same SNP after 45 min. In addition to its exceedingly high specificity, quantification of 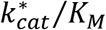 could potentially be automated and parallel measurements at various dilutions can potentially be completed faster than any endpoint fluorescence experiments since the relevant initial cleavage rates can be measured within about 5 min. A limitation of this assay approach is, of course, that it requires knowledge of the target concentration.

**Fig. 4.**
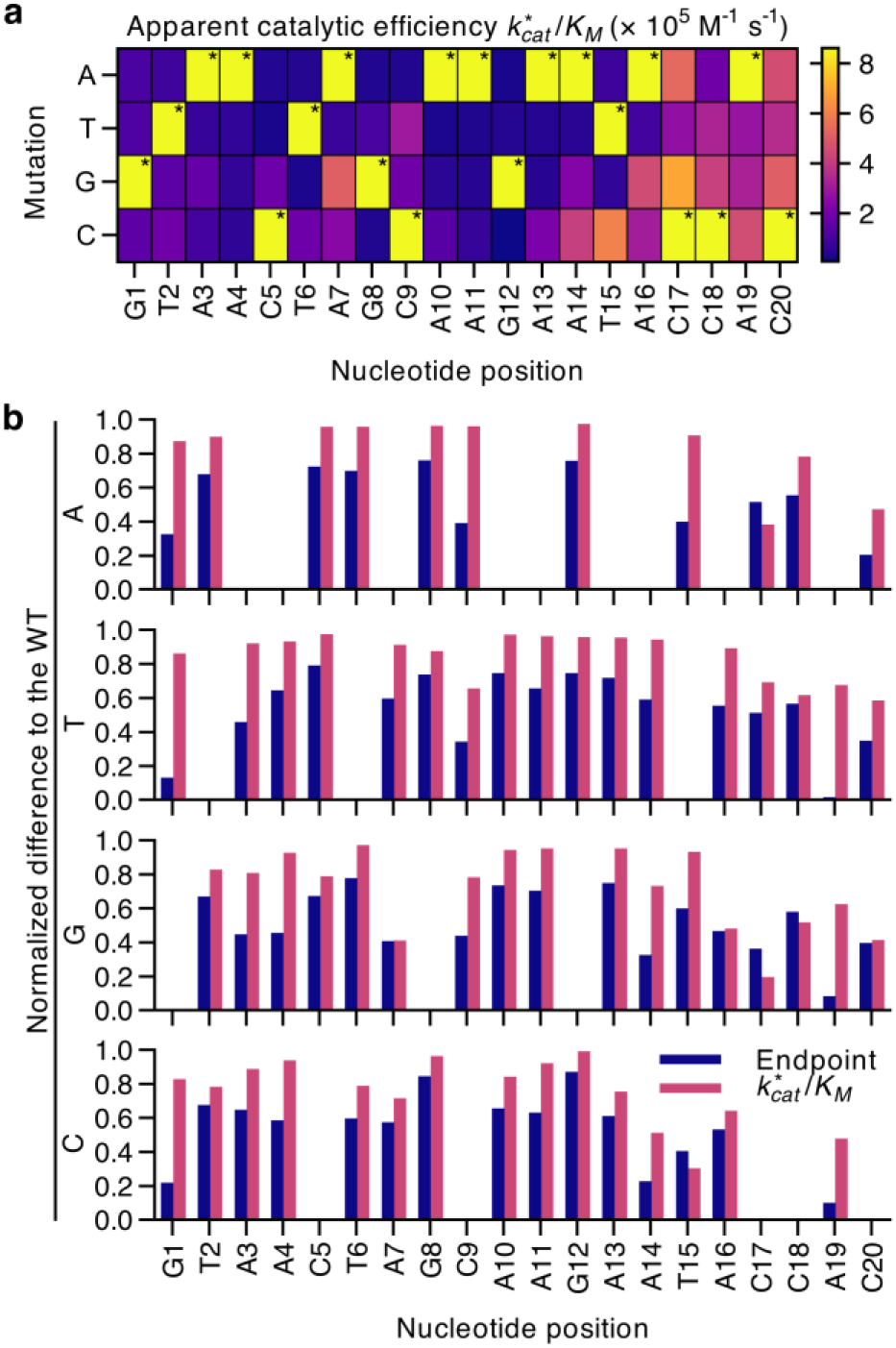
Catalytic efficiencies of WT and all SNPs and their normalized comparison to end-point detection. **a** Measured catalytic efficiency 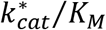 for WT and all SNPs, as determined from analysis of the Michaelis-Menten curves. Asterisks denote the cells which correspond to WT. For all cases, the 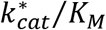 of the WT was consistently larger than all SNPs. **b** Normalized difference to the WT endpoint fluorescence after 45 min (in blue) or apparent catalytic efficiency 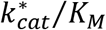 (in pink). Each plot row corresponds to mutations to the base indicated in the ordinate. Higher values correspond to a greater ability to discriminate among WT and SNP targets. 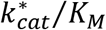 was a better discriminator of WT versus SNP than endpoint fluorescence for 57 SNPs and worse only for T15C, C17A, C17G, and C18G. Blank entries (no bars) indicate the WT.

As in the case of the *K_M_* data, we found that SNPs at nucleotide positions near the 3’ end (#17 to 20) had 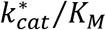 values closer to that of the WT. To further evaluate the effect of SNP position on either endpoint- or catalytic efficiency-based differentiation of WT, we calculated their normalized difference to the WT (**Fig. 4b**). We term this difference *D* and calculate mathematically as

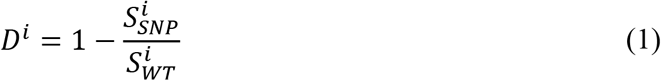

where the superscript *i* denotes either detection by endpoint fluorescence after 45 min (*i* = *EF*) or detection via quantification of 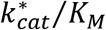 (*i* = *CE*), *S* is the relevant signal, and the subscript *SNP* is a variable to designate a particular SNP or WT. For example, 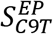 corresponds to the endpoint fluorescence of SNP C9T while 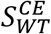 corresponds to the 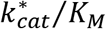 of the WT. Interestingly, we found that *D^CE^* was greater than *D^EF^*, indicating an improved degree of discrimination, for all associated SNPs except T15C, C17A, C17G, and C18G. Also, we further observed that, for a fixed mutation base (say, mutations to A), *D_CE_* mostly decreased with increasing nucleotide position (nearer to the 3’ end) while *D_EF_* was greatest towards the middle nucleotide positions.

Next, we used an experimentally validated Michaelis-Menten model^17^ to show scenarios wherein target concentration is not known and the studied SNPs are nearly indistinguishable from WT using EP detection. Specifically, we predicted progress curves for the measured 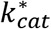 and *K_M_* values of WT and all mutants (*c.f*. **Figs. 2** and **3**) and equal values of substrate concentration. For the WT, we fixed target concentration at 1 nM. For each mutant case, we then varied target concentration to predict the progress curve which minimized the mean-squared error (MSE) between the WT and SNP *trans*-cleavage progress curves (**Fig. 5a**). This simple use of the model demonstrates how variations in target concentration can result in SNP progress curves that are nearly indistinguishable from the WT. In turn, the magnitude of the WT-to-SNP cleaved reporter concentration difference was less than 20 nM, equivalent to 10% of the total reporter concentration (inset of **Fig. 5a**). We also found that faster-than-WT SNP progress curves were most different from the WT around 0.5 h after the experiment commenced as opposed to 2 h for slower-than-WT progress curves. We hypothesize the latter trend may be due to the underlying physics of the Michaelis-Menten kinetics model.

**Fig. 5.**
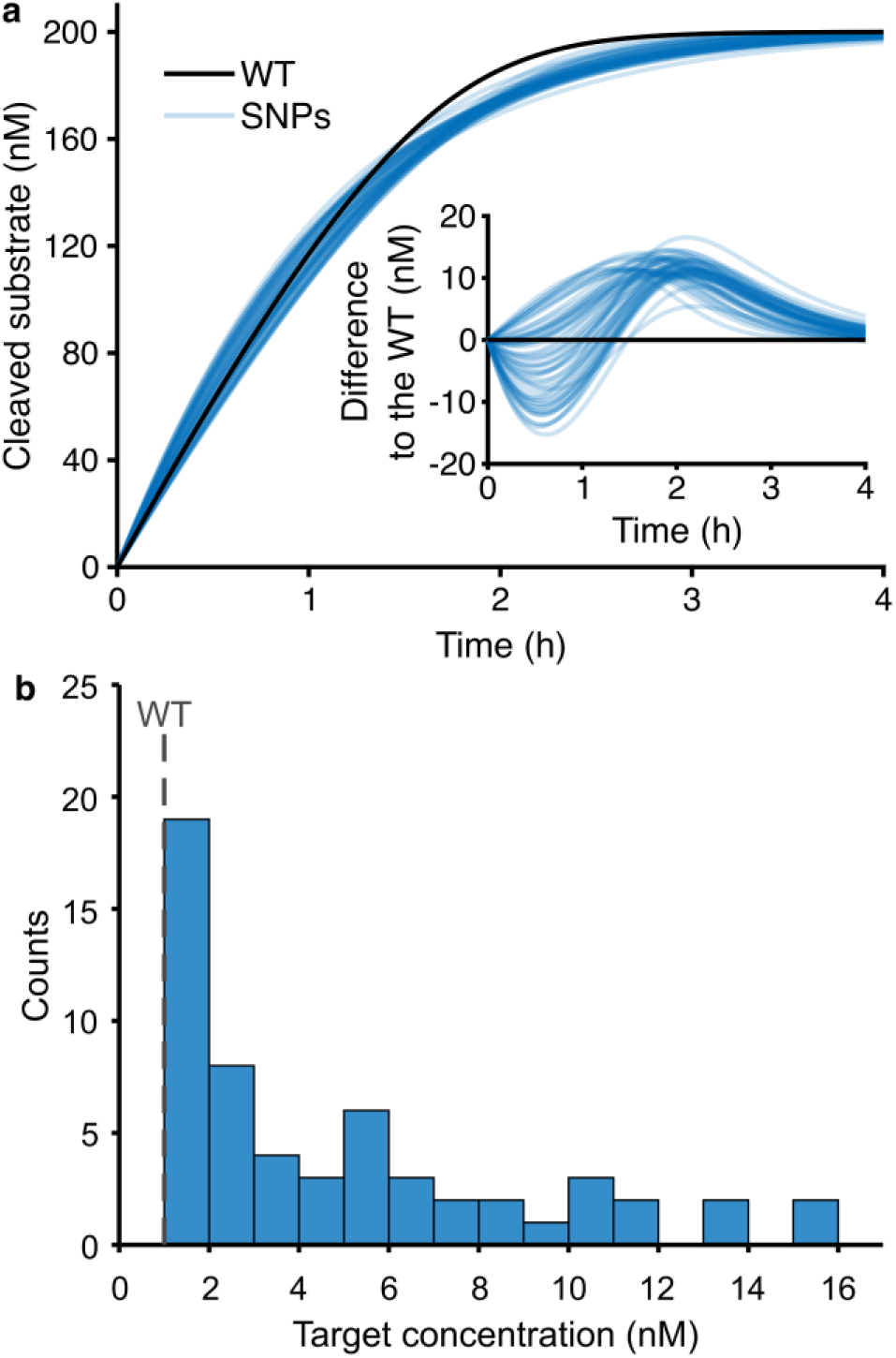
Example predicted scenarios wherein endpoint fluorescence is unlikely to differentiate the signal from SNP targets relative to WT. All simulations used kinetic rate parameters measured here (**Figs. 2** and **3**). **a** Predicted *trans*-cleavage progress curves (of cleaved substrate concentration) versus time. Each of the 60 blue lines was generated by fixing the corresponding kinetic parameters of a SNP and then varying SNP target concentration to minimize the mean-squared error relative to WT. The inset shows the cleaved reporter concentration difference (WT minus SNP, in molar units) versus time for the same progress curves. Curves in both plots are slightly transparent to facilitate visualization. **b** Histogram of the computed target concentrations for each of the 60 SNPs of **a** required to obtain a progress curve similar to the WT (the WT concentration was 1 nM).

A histogram of the computed SNP target concentrations which gave minimum MSE relative to WT showed that 19 out of the 60 computed SNP concentrations were less than 2-fold different than the WT (**Fig. 5b**). Further, 34 SNPs (representing over half of all studied SNPs) had target concentrations within 5-fold of the WT. These observations highlight the notion that precise knowledge of SNP target concentration would be needed to distinguish SNPs from WT.

In conclusion, we here proposed fundamental Michaelis-Menten parameters as a time- and target concentration-insensitive modality for SNP differentiation. We further showed that our proposed method yielded greater dynamic range than endpoint fluorescence and could potentially provide faster detection (since initial reaction velocities can be measured within 5 min). Finally, we used an experimentally validated model to show how SNP progress curves can be practically indistinguishable from WT when SNP concentration is not precisely known.

## METHODS

### Complexing and activating Cas12-gRNA

LbCas12a was purchased from New England Biolabs (MA, USA) at a concentration of 100 μM. RNA oligonucleotides were purchased from GeneLink (FL, USA) and resuspended to 100 μM in RNA reconstitution buffer (GeneLink). ssDNA WT and SNP targets were purchased from Elim Biopharmaceuticals Inc. (CA, USA) at 100 μM. A complete list of oligos is provided in the Supplementary Information (**Table S2**). Our complexing and activating protocol were as follows. We first prepared 100 nM solutions of the Cas12-gRNA complex. This was achieved by incubating a mixture of 1 μM Cas12 with at least 5-fold excess of synthetic gRNA at 37 °C for 30 min on a hot plate. Cas12-gRNA complexes were then activated for *trans*-cleavage activity after incubation with either WT target or one of the 61 SNPs. Specifically, Cas12-gRNA complexes were incubated with at least 5-fold excess of the synthetic ssDNA at 37 °C for 30 min on a hot plate. The latter step yielded a solution with an activated Cas12 concentration of 50 nM.

### Trans-cleavage kinetics experiments

We performed the *trans-cleavage* kinetics assay using 2 nM of activated Cas12 and varied ssDNA reporter concentrations of 13, 25, 50, 100, 200, 400, 800, and 1600 nM. Three replicates were taken for each concentration. The final volume of each reaction was 30 μL. All reactions were buffered in 1× NEBuffer 2.1 (composed of 50 mM NaCl, 10 mM Tris-HCl, 10 mM MgCl2, and 100 μg mL^-1^ bovine serum albumin at pH = 7.9). All *trans*-cleavage reactions were run at 37 °C. Cas12 kinetics data was measured using a 7500 Fast Real-Time PCR system (Applied Biosystems, CA, USA), and fluorescence was measured every 60 s.

### Analysis of progress curves

All *trans*-cleavage and calibration data points were first background and flatfield corrected. We found that our thermal cycler (7500 Fast Real-Time PCR system, Applied Biosystems) showed significant well-to-well (position-to-position) differences in fluorescence signals for equal concentrations of fluorophores. Hence, by flatfield correction, we here mean a correction for (repeatable and measurable) non-uniform response of the instrument across wells. We term the discrete array of fluorescence values an “image” of the wells. To obtain a background image, wells were filled with 1× NEBuffer 2.1. For the flat field image, wells were filled with 1.6 μM fluorescein. The correction is then as follows.

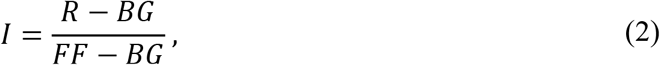

where *I* is the corrected image, *R* is the raw image, *BG* is the background image, and *FF* is the flatfield image. Note that in **Eq. 3**, only *I* and *R* contain time series data (since *BG* and *FF* were averaged in time).

A fluorescence calibration curve was then used to convert fluorescence values (in AU) to molar concentrations (in nM).^19,20^ To this end, two nonlinear fits were performed on calibration data, one each for cleaved and uncleaved reporters (**Fig. S1**). The exponential law fit has the form

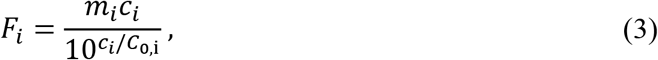

where the subscript *i* denotes the uncleaved (= *ucl*) or cleaved (= *cl*) data, *F* is the (background and flatfield) corrected fluorescence, *m* is the slope of the linear regime of the cleaved or uncleaved reporter calibration curve, *c* is the reporter concentration, and *C*_0_ is the value that is varied to obtain the best fit. Once *C*_0,*uct*_ and *C*_0,*cl*_ were obtained, the conversion of fluorescence values to molar concentration of cleaved reporters followed the equation

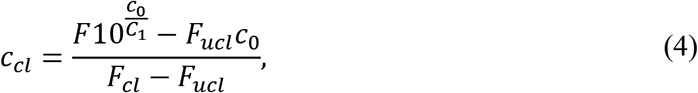

where

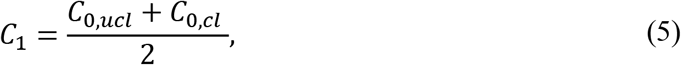

*F* is the measured fluorescence during the *trans*-cleavage experiment, *F_ucl_* is the slope of the linear regime of the uncleaved fluorescence intensity versus reporter concentration best fit line, *F_cl_* is the slope of the linear regime of the cleaved fluorescence intensity versus reporter concentration best fit line, and *c*_0_ is the total reporter concentration for each experiment.

The initial reaction velocities (in nM s^-1^) for Michaelis-Menten analyses were obtained using a linear fit to the first ~300 s of the *trans*-cleavage progress curves. The reaction velocities versus reporter concentration data were fitted to the Michaelis-Menten equation using GraphPad Prism 9 (GraphPad Software, CA, USA) to obtain *V_max_* and *K_M_*.

## ACKNOWLEDGEMENTS

The authors gratefully acknowledge funding from Stanford University’s Interdisciplinary Biosciences Institute (Bio-X) and from Stanford’s School of Engineering COVID-19 Research and Assistance Fund. C.B. acknowledges support from the France-Stanford Center. D.A.H. is supported by a National Science Foundation Graduate Research Fellowship. A.R. acknowledges support from the Bio-X Bowes Fellowship of Stanford University.

## Notes

### Competing Interest Statement

The authors have declared no competing interest.

