## Supplementary Information for "Detection and discrimination of single nucleotide polymorphisms by quantification of CRISPR-Cas catalytic efficiency"

**Contents**

S1 Example of C9T endpoint fluorescence
S2 DNA reporter calibration curves
S3 Michaelis-Menten fits for all SNPs
S4 Michaelis-Menten fits for combined WT and SNP targets
S5 Michaelis-Menten kinetic parameters with confidence intervals
S6 List of oligos used in this work

### S1 Example of C9T endpoint fluorescence

In this section, we present predictions of the *trans*-cleavage activity of WT and C9T (**Fig. S1**) based on measured Michaelis-Menten parameters (**Table S1**) for the experimental conditions presented in the main manuscript (**Fig. 1d**). The predictions presented here were generated using an experimentally validated Michaelis-Menten model.[1]

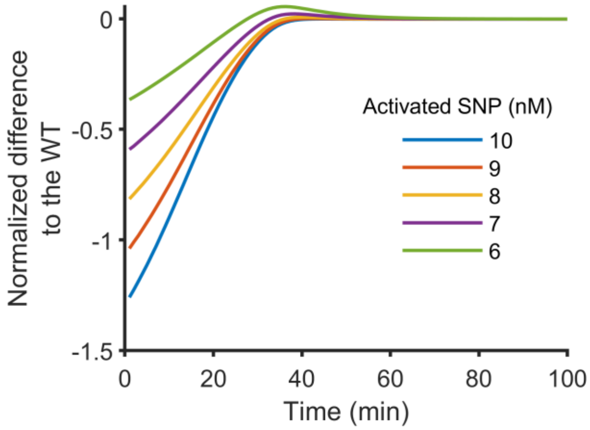


**Fig. S1 SNP C9T has a high *trans*-activated turnover rate**. Difference to WT fluorescence signal (normalized by WT signal) versus time for various *trans*-activated concentrations of C9T. Plot was generated in MATLAB (R2021b, Mathworks, USA) using an experimentally validated numerical model of Michaelis-Menten kinetics. The kinetic parameters ($k_{cat}$ and $K_M$) for the WT and C9T were extracted from Michaelis-Menten curves for those same targets (**Table S1**). The WT-activated enzyme and initial substrate concentration were respectively 10 and 800 nM (these are the same experimental conditions as **Fig. 1d**). Interestingly, the low SNP concentration conditions (= 6 and 7 nM) resulted in substrate cleavage initially faster and then slower than WT. All conditions resulted in no difference to WT (when all substrate was fully cleaved) for times greater than 80 min.

### S2 DNA reporter calibration curves

**Fig. S2** shows the calibration curves for the cleaved and uncleaved ssDNA reporters used in this work. The calibration curves were used to quantify cleavage activity in molar units.[2]

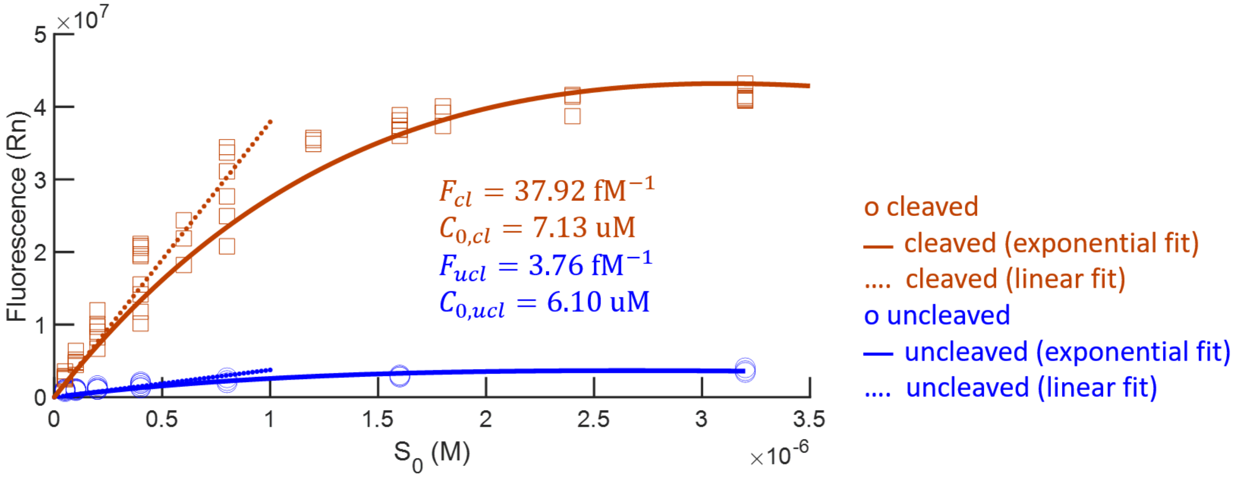


**Fig. S2 RNA reporter calibration curves**. Measured fluorescence versus concentration of uncleaved and cleaved FAM-BHQ DNA reporters. Lines (dashed line) and power law curves (solid lines) of best fit to the experimental data were obtained by linear regression. Data was taken in a 7500 Fast Real-Time PCR system (Applied Biosystems, CA, USA). Significant inner filter effect leads to the non-linearity,[3] and this is accounted for by the exponential law (see **Methods** in the main manuscript).

### S3 Michaelis-Menten fits for all SNPs

This section presents Michaelis-Menten fits to the initial reaction for all SNPs at all positions (**Figs. S3-S5**). These data comprise measurements across 60 SNPs over 20 nucleotide positions.

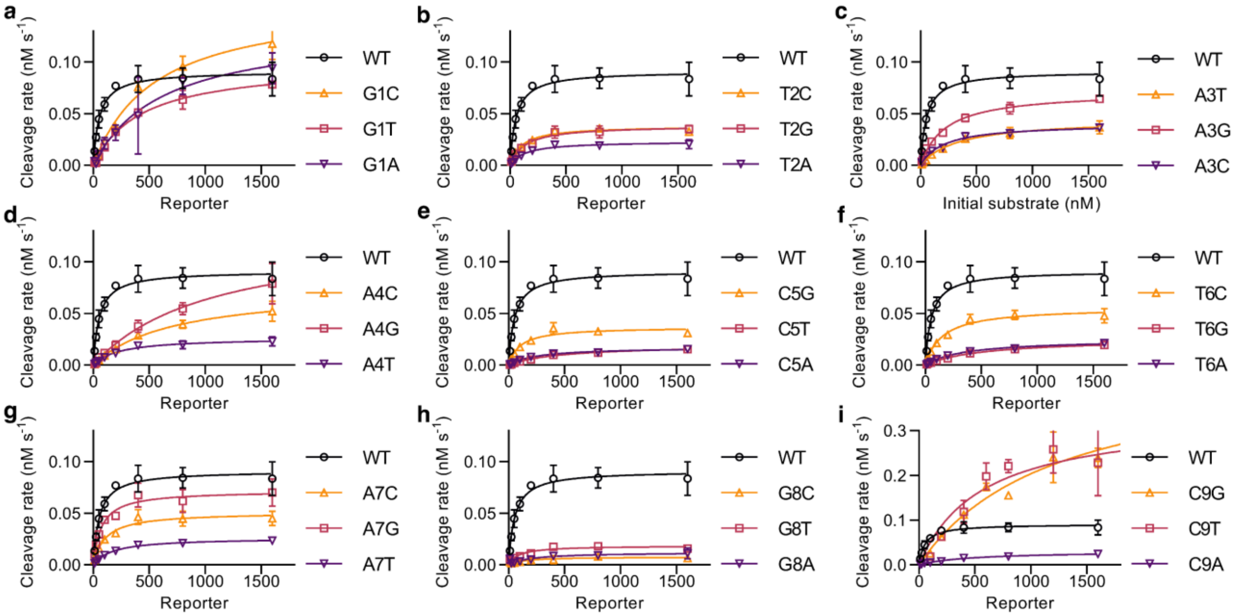


**Fig. S3 LbCas12 Michaelis-Menten fits for all SNPs in nucleotide positions #1-9. a** Cleavage rate versus initial substrate (uncleaved reporter) concentration for WT and SNPs at position #1. Shown together with the data are fits to the Michaelis-Menten equation. **b-i** Show same data as **a** for WT and, respectively, SNPs in positions #2-9. The Cas12-gRNA complex concentration for all experiments was 2 nM.

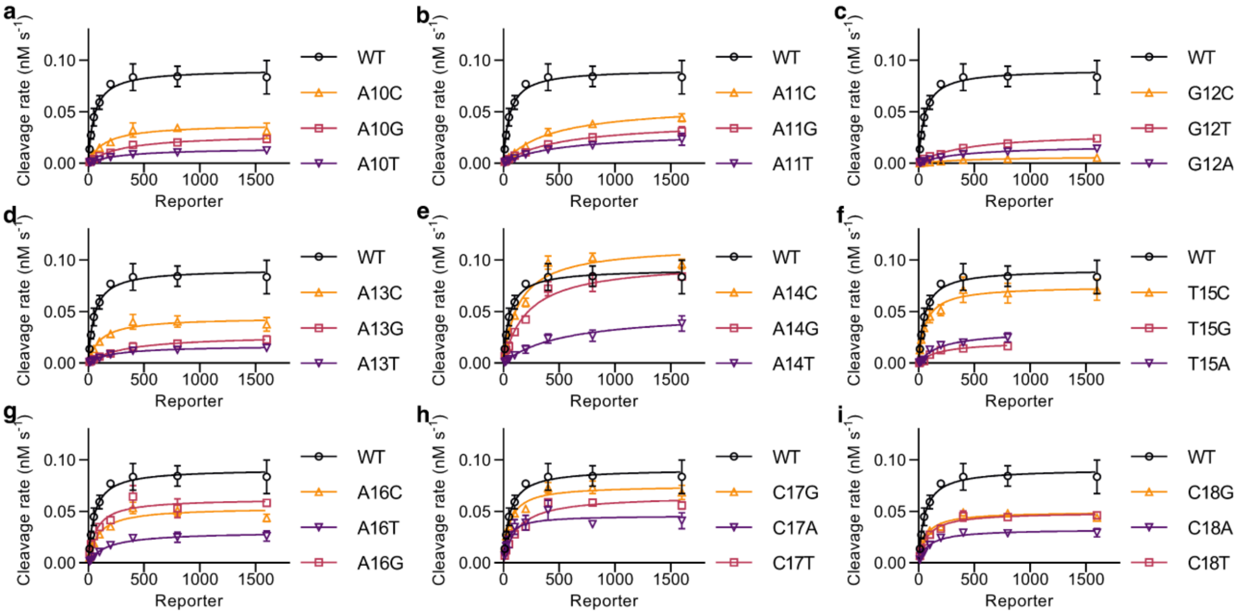


**Fig. S4 LbCas12 Michaelis-Menten fits for all SNPs in nucleotide positions #10-18. a** Cleavage rate versus initial substrate (uncleaved reporter) concentration for WT and SNPs at position #10. Also shown are fits to the Michaelis-Menten equation. **b-i** Show same data as **a** for WT and, respectively, SNPs in positions #11-18. The Cas12-gRNA complex concentration for all experiments was 2 nM.

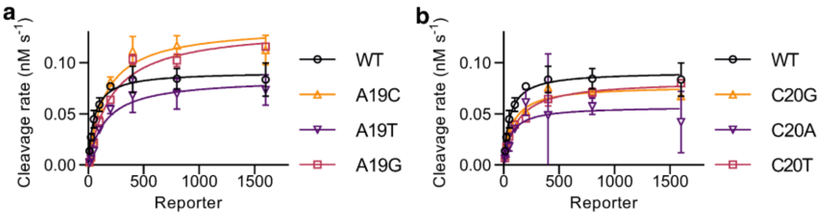


**Fig. S5 LbCas12 Michaelis-Menten fits for all SNPs in nucleotide positions #19 and 20.** Cleavage rate versus initial substrate (uncleaved reporter) concentration for WT and SNPs at position #19. Also shown are fits to the Michaelis-Menten equation. **b** Shows same data as **a** for WT and SNPs in position #20. The Cas12-gRNA complex concentration for all experiments was 2 nM.

### S4 Michaelis-Menten fits for combined WT and SNP targets

This section presents experimental results wherein Cas12-gRNA complex was activated by combined WT and SNP targets (**Fig. S6**).

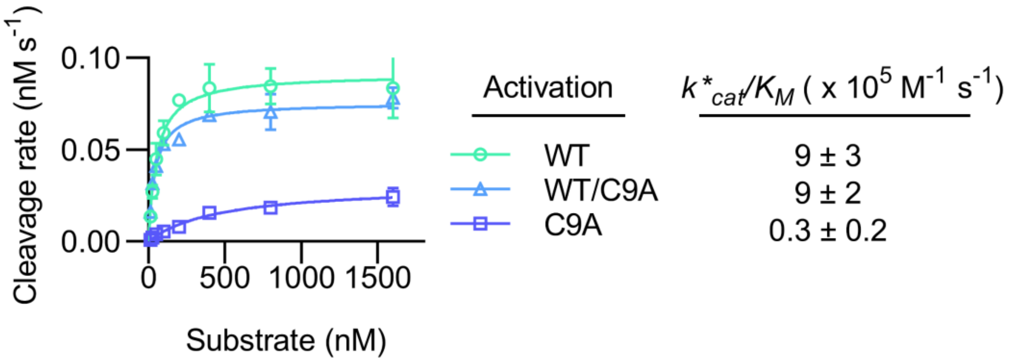


**Fig. S6 Michaelis-Menten fits for combined WT and SNP experiments.** Cleavage rate versus initial substrate (uncleaved reporter) concentration for WT, SNP C9A, and the 50/50 WT/C9A mix. Also shown are fits to the Michaelis-Menten equation. The Cas12-gRNA complex concentration for all experiments was 2 nM. Target concentrations for WT, C9A, and WT/C9A mix were respectively 5-fold excess, 5-fold excess, and 2.5-fold excess for both WT and C9A targets in the mix. Results suggest that almost all Cas12-gRNA complexes were trans-activated by WT targets due to their higher affinity.

### S5 Michaelis-Menten kinetic parameters with confidence intervals

**Table S1** shows confidence intervals for the measured kinetic parameters of the WT and all SNPs. Confidence intervals are estimates obtained from GraphPad Prism 9 (GraphPad Software, CA, USA) using the asymmetrical (profile-likelihood) confidence interval option.

**Table S1 Summary of experimentally measured kinetic parameters.** This table reports the kinetic parameters and 95% confidence intervals on $k_{cat}^{*}$ and $K_M$ measurements.

| Target | Best fit | | 95% confidence intervals | |
|---|---|---|---|---|
| | $k_{cat}^{*}$ ($s^{-1}$) | $K_M$ (nM) | $k_{cat}^{*}$ ($s^{-1}$) | $K_M$ (nM) |
| WT | 0.046 | 53 | 0.042 to 0.049 | 39 to 72 |
| G1A | 0.067 | 620 | 0.05 to 0.098 | 310 to 1300 |
| G1T | 0.05 | 430 | 0.045 to 0.055 | 340 to 540 |
| G1C | 0.08 | 540 | 0.071 to 0.091 | 410 to 730 |
| T2A | 0.012 | 140 | 0.01 to 0.013 | 93 to 200 |
| T2G | 0.019 | 130 | 0.018 to 0.021 | 95 to 180 |
| T2C | 0.019 | 100 | 0.018 to 0.02 | 80 to 130 |
| A3T | 0.022 | 340 | 0.02 to 0.025 | 250 to 460 |
| A3G | 0.036 | 220 | 0.034 to 0.038 | 180 to 270 |
| A3C | 0.02 | 210 | 0.019 to 0.022 | 160 to 270 |
| A4T | 0.013 | 230 | 0.012 to 0.015 | 160 to 330 |
| A4G | 0.065 | 1100 | 0.051 to 0.092 | 640 to 1900 |
| A4C | 0.038 | 720 | 0.033 to 0.045 | 510 to 1000 |
| C5A | 0.009 | 260 | 0.008 to 0.01 | 190 to 340 |
| C5T | 0.01 | 480 | 0.009 to 0.011 | 390 to 580 |
| C5G | 0.018 | 100 | 0.017 to 0.02 | 73 to 140 |
| T6A | 0.013 | 360 | 0.011 to 0.014 | 270 to 480 |
| T6G | 0.013 | 530 | 0.012 to 0.015 | 410 to 700 |
| T6C | 0.028 | 150 | 0.026 to 0.03 | 120 to 200 |
| A7T | 0.013 | 180 | 0.012 to 0.014 | 140 to 220 |
| A7G | 0.036 | 71 | 0.033 to 0.039 | 49 to 100 |
| A7C | 0.025 | 110 | 0.023 to 0.028 | 74 to 150 |
| G8A | 0.006 | 200 | 0.004 to 0.01 | 51 to 710 |
| G8T | 0.009 | 89 | 0.008 to 0.01 | 55 to 140 |
| G8C | 0.004 | 130 | 0.003 to 0.005 | 46 to 320 |
| C9A | 0.015 | 470 | 0.013 to 0.018 | 320 to 690 |
| C9T | 0.17 | 570 | 0.14 to 0.21 | 320 to 990 |
| C9G | 0.23 | 1200 | 0.19 to 0.28 | 840 to 1800 |
| A10T | 0.007 | 310 | 0.007 to 0.008 | 220 to 430 |
| A10G | 0.014 | 300 | 0.013 to 0.015 | 250 to 350 |
| A10C | 0.019 | 140 | 0.017 to 0.021 | 96 to 210 |
| A11T | 0.015 | 520 | 0.013 to 0.019 | 330 to 860 |
| A11G | 0.021 | 510 | 0.018 to 0.023 | 380 to 700 |
| A11C | 0.029 | 420 | 0.026 to 0.031 | 340 to 530 |
| G12A | 0.009 | 410 | 0.008 to 0.01 | 290 to 600 |
| G12T | 0.015 | 410 | 0.013 to 0.017 | 320 to 550 |

| | | | | |
|---|---|---|---|---|
| G12C | 0.004 | 550 | 0.003 to 0.005 | 330 to 950 |
| A13T | 0.009 | 220 | 0.008 to 0.01 | 160 to 310 |
| A13G | 0.014 | 340 | 0.012 to 0.016 | 230 to 490 |
| A13C | 0.022 | 110 | 0.02 to 0.024 | 74 to 150 |
| A14T | 0.025 | 540 | 0.02 to 0.032 | 330 to 910 |
| A14G | 0.049 | 220 | 0.045 to 0.054 | 170 to 280 |
| A14C | 0.057 | 140 | 0.053 to 0.062 | 107 to 180 |
| T15A | 0.016 | 200 | 0.013 to 0.019 | 130 to 310 |
| T15G | 0.01 | 180 | 0.009 to 0.013 | 120 to 290 |
| T15C | 0.037 | 63 | 0.034 to 0.041 | 44 to 88 |
| A16T | 0.015 | 170 | 0.014 to 0.017 | 120 to 240 |
| A16G | 0.031 | 70 | 0.028 to 0.034 | 47 to 103 |
| A16C | 0.027 | 88 | 0.024 to 0.03 | 61 to 120 |
| C17A | 0.023 | 43 | 0.02 to 0.026 | 25 to 72 |
| C17T | 0.033 | 120 | 0.03 to 0.035 | 94 to 160 |
| C17G | 0.038 | 54 | 0.035 to 0.04 | 42 to 70 |
| C18A | 0.016 | 89 | 0.015 to 0.017 | 70 to 110 |
| C18T | 0.025 | 75 | 0.023 to 0.026 | 60 to 94 |
| C18G | 0.025 | 60 | 0.023 to 0.026 | 47 to 77 |
| A19T | 0.043 | 150 | 0.037 to 0.049 | 99 to 240 |
| A19G | 0.067 | 210 | 0.063 to 0.073 | 170 to 260 |
| A19C | 0.068 | 150 | 0.062 to 0.075 | 110 to 210 |
| C20A | 0.029 | 63 | 0.02 to 0.039 | 15 to 210 |
| C20T | 0.041 | 120 | 0.039 to 0.044 | 94 to 140 |
| C20G | 0.039 | 77 | 0.036 to 0.041 | 61 to 98 |

**S6 List of oligos used in this work**

**Table S2** lists all oligos used in this work, including the gRNA as well as WT and SNP targets.

**Table S2. List of gRNAs used in this work.** Vendors for all oligos and the reconstitution procedures can be found in the **Methods** section of the main manuscript.

| **Name** | **Type** | **Sequence (5'-3')** | **Ref** |
|---|---|---|---|
| gRNA | ssRNA | UAA UUU CUA CUA AGU GUA GAU GUG GUA UUC UUG CUA GUU AC | 1 |
| WT* | ssDNA | G TAA CTA GCA AGA ATA CCA C | |
| Rep | ssDNA | [5-Fam] TTA TTA TT [3-BHQ-1] | |

* Note all SNPs were generated from this WT sequence.
